# Conformational Flexibility and Local Frustration in the Functional States of the SARS-CoV-2 Spike B.1.1.7 and B.1.351 Variants : Mutation-Induced Allosteric Modulation Mechanism of Functional Dynamics and Protein Stability

**DOI:** 10.1101/2021.12.22.473892

**Authors:** Gennady Verkhivker

## Abstract

The experimental and computational studies of the SARS-CoV-2 spike protein variants revealed an important role of the D614G mutation that is shared across variants of concern(VOCs), linking the effect of this mutation with the enhanced virus infectivity and transmissibility. The recent structural and biophysical studies characterized the closed and open states of the B.1.1.7 (B.1.1.7) and B.1.351 (Beta) spike variants allowing for a more detailed atomistic characterization of the conformational landscapes and functional changes. In this study, we employed coarse-grained simulations of the SARS-CoV-2 spike variant trimers together with the ensemble-based mutational frustration analysis to characterize the dynamics signatures of the conformational landscapes. By combining the local frustration analysis of the conformational ensembles with collective dynamics and residue-based mutational scanning of protein stability, we determine protein stability hotspots and identify potential energetic drivers favoring the receptor-accessible open spike states for the B.1.1.7 and B.1.351 spike variants. Through mutational scanning of protein stability changes we quantify mutational adaptability of the S-G614, S-B.1.1.7 and S-B.1.351 variants in different functional forms. Using this analysis, we found a significant conformational and mutational plasticity of the open states for all studied variants. The results of this study suggest that modulation of the energetic frustration at the inter-protomer interfaces can serve as a mechanism for allosteric couplings between mutational sites, the inter-protomer hinges of functional motions and motions of the receptor-binding domain required for binding of the host cell receptor. The proposed mechanism of mutation-induced energetic frustration may result in the greater adaptability and the emergence of multiple conformational substates in the open form. This study also suggested functional relationships between mutation-induced modulation of protein dynamics, local frustration and allosteric regulation of the SARS-CoV-2 spike protein.

## 1. Introduction

SARS-CoV-2 infection is transmitted when the viral spike (S) glycoprotein binds to the host cell receptor ACE2, leading to the entry of S protein into host cells and membrane fusion [1,2]. The full-length SARS-CoV-2 S protein consists of amino (N)-terminal S1 subunit and carboxyl (C)-terminal S2 subunit where S1 is involved in the interactions with the host receptor and includes an N-terminal domain (NTD), the receptor-binding domain (RBD), and two structurally conserved subdomains (SD1 and SD2). Structural and biochemical studies established that the mechanism of virus infection may involve conformational transitions between distinct functional forms of the SARS-CoV-2 S protein in which the RBDs continuously switch between “down” and “up” positions [3–10]. The cryo-EM structure of the SARS-CoV-2 S trimer revealed a spectrum of closed states that included a structurally rigid closed form and more dynamic closed states preceding a transition to the fully open S conformation [5]. Protein engineering and structural studies also showed that specific disulfide bonds and proline mutations can modulate stability of the SARS-CoV-2 S trimer [6] and lead to the thermodynamic shifts between the closed and open forms [7–9]. Dynamic structural changes that accompany SARS-CoV-2 S binding with the ACE2 host receptor were described in cryo-EM experiments showing a cascade of conformational transitions from a compact closed form weakened after furin cleavage to the partially open states and subsequently the ACE2-bound open form thus priming the S protein for fusion [10]. The cryo-EM and tomography examined conformational flexibility and distribution of S trimers in situ on the virion surface [11] showing that spontaneous conformational changes and population shifts between different functional states can be maintained in different biological environments, reflecting the intrinsic properties of conformational landscapes for SARS-CoV-2 S trimers. Single-molecule Fluorescence (Förster) Resonance Energy Transfer (smFRET) studies of SARS-CoV-2 S trimer on virus particles revealed a sequence of conformational transitions from the closed state to the receptor-bound open state suggesting that mechanisms of conformational selection and receptor-induced structural adaptation can work synchronously and showing that SARS-CoV-2 neutralization can be achieved by antibodies that either directly compete with the ACE2 receptor binding or exert their effect by allosterically stabilizing the S protein in its RBD-down conformation [12]. The expanding body of structural and biochemical studies of the SARS-CoV-2 S complexes with different classes of potent antibodies targeting distinct binding epitopes of the S-RBD as well as various antibody cocktails and combinations have revealed multiple conformation-dependent epitopes, highlighting the link between conformational plasticity and adaptability of S proteins and capacity for eliciting specific binding and broad neutralization responses [13–36]. Deep mutagenesis scanning of antibody-escaping mutations showed that the escape mutations cluster in several RBD regions and can be constrained with respect to their effects on expression of properly folded RBD and ACE2 binding [37–42]. SARS-CoV-2 S mutants with the enhanced infectivity profile including D614G mutational variant have attracted an enormous attention in the scientific community following the evidence of the mutation enrichment via epidemiological surveillance, resulting in proliferation of experimental data and a considerable variety of the proposed mechanisms explaining functional observations [43–45]. The latest biochemical studies provided evidence of a phenotypic advantage and the enhanced infectivity conferred by the D614G mutation [46]. The initial structural studies showed that the D614G mutation can act by shifting the population of the SARS-CoV-2 S trimer from the closed form (53% of the equilibrium) in the native spike protein to a widely-open topology of the “up” protomers in the D614G mutant with 36% of the population adopting a single open protomer, 39% with two open protomers and 20% with all three protomers in the open conformation [47]. The cryo-EM structures of the S-D614 and S-G614 ectodomains showed the increased population of the 1-RBD-up open form as compared to the closed state in the S-GSAS/D614G structure [48]. The electron microscopy analysis also revealed the higher 84% percentage of the 1-up RBD conformation in the S-G614 protein [49]. Functional studied showed that the S-G614 mutant exhibited the greater infectivity than the S-D614 protein which was attributed to the greater stability of the S-G614 mutant and leading to the reduced S1 subdomain shedding [50]. The increased stability of the D614G mutant was inferred from the recent cryo-EM structures of a full-length unmodified S-G614 trimer that can reversibly adopt an all-down closed state and 1 RBD-up open conformation [51].

B.1.1.7 variant of the SARS-CoV-2, a descendant of the D614G lineage, has originated in UK and spread to 62 countries showing the increased transmissibility. 8 of the 17 mutations observed in this variant (Δ69-70 deletion, Δ144 deletion, N501Y, A570D, P681H, T716I, S982A, D1118H) are accumulated in the S protein, featuring most prominently N501Y mutation that can increase binding affinity with ACE2 while eliciting immune escape and reduced neutralization of RBD-targeting antibodies [52–54]. SARS-CoV-2-S pseudo-viruses bearing either the reference strain or the B.1.1.7 lineage spike protein with sera of 40 participants who were vaccinated with the mRNA-based vaccine BNT162b2 showed largely preserved neutralization, indicating that the B.1.1.7 lineage will not escape BNT162b2-mediated protection [55]. SARS-CoV-2 B.1.351 variant was first detected in South Africa is characterized by 21 mutations with nine mutations (L18F, D80A, D215G, R246I, K417N, E484K, N501Y, D614G, and A701V) in the spike protein, of which three mutations (K417N, E484K, and N501Y) are located in the RBD and increase the binding affinity for the ACE receptors and induce significant immune escape [56,57]. The recent data demonstrate reduced but still significant neutralization against the full B.1.351 variant following mRNA-1273 vaccination [58]. Functional and structural studies explored how B.1.1.7 (B.1.1.7), B.1.351 (beta), P1 (gamma) and B.1.1.427/B.1.429 (epsilon) variants in the SARS-CoV-2 S protein affect conformational landscapes of the S protein and the ability to evade host immunity and incur resistance to antibodies [59–69]. These studies revealed subtle structural and functional impact of mutations that can modulate dynamics and stability of the closed and open forms, increase binding to the human receptor ACE2, and confer resistance to neutralizing antibodies. Structural determination of the SARS-CoV-2 S S-B.1.1.7 trimer showed preferences for the RBD-up conformation. The FPPR (residues 828 to 853) and 630 loop (residues 620 to 640) modulate the stability and structural rearrangements of the S protein [70]. Based on structural evidence, it was proposed that B.1.1.7 mutations A570D and S982A can induce global movement of the CTD1, thereby relaxing the FPPR and 630 loop, which may increase the frequency with which the S trimer samples the RBD-up conformation. The overall structures of the S-B.1.351 and S-G614 trimers were similar for the corresponding states, with mutations K417N, E484K, and N501Y resulting in only minor structural rearrangements [70]. The cryo-EM structures of the S protein of B.1.1.7 variant identified four distinct conformational states with three of the conformational classes corresponded to a 1 RBD-up and one to 2 RBD-up conformation [71]. This study indicated that B.1.1.7 mutations can modulate dynamics of the open S conformations without introducing substantial structural changes as compared to the S-G614 structures. Importantly, it was proposed that A570D can serve as a molecular switch that could modulate the opening and closing of the RBD [71]. In this mechanism, the switching of the A570D-mediated salt bridges may serve as a pedal-bin-like device to modulate the RBD motion. A total of 38 cryo-EM structures of the S-B.1.351, S-P.1 (Gamma), S-B.1.617.2 (Delta) and S-B.1.617.1 (Kappa) variants in different functional states with and without its receptor ACE2 have been recently reported showing a diverse and vast conformational landscape for each of the variants [72]. The structures of these S variants confirmed the broad spectrum of intrinsic RBD-up propensities that are inherently present in the apo states, with different numbers of RBD-up and multiple substates. In particular, S-B.1.351 variant displayed a highly populated fully open state with all three RBDs in the up conformation in addition to significant fractions of the partially open states (1 RBD-up or 2 RBD-up) [72]. Importantly, among all the S variants only S-B.1.351 exhibited a fully open 3 RBD-up conformation that was not observed in related structural studies [63,70,71] and that resembles structure of the fully open S-G614 variant [47]. These structural studies suggested that the increased conformational diversity and particularly greater variability of the open states can allow S-Beta, S-Gamma and S-Delta variants to adapt their responses and provide mechanisms for escape immunity from common vaccines and different classes of monoclonal antibodies. Biophysical studies using microscale thermophoresis established that SARS-CoV-2 B.1.1.7 and B.1.351 spike variants bind human ACE2 with increased affinity [73]. According to this study, the enhanced affinity can mediate increased infectivity by lowering the effective concentration of virions required for cell entry which would result in highly transmissible SARS-CoV-2 variants. Using surface plasmon resonance (SPR), the effects of individual RBD mutations and combinations found in S-Alpha, S-Beta and S-Gamma variants on thermodynamics and kinetics of ACE2 binding [74]. It was found that most of these mutations increased the affinity of the RBD/ACE2 interaction with the exception of K417N/T which decreased the affinity. Collectively, the body of structural and biochemical studies revealing the emerging conformational plasticity of the S variants suggested that coronaviruses have broad potential to tolerate both sequence and structure changes without substantial loss of function [75]. The detection of common mutational changes D614G, E484K, N501Y and K417N shared among major circulating variants B.1.1.7, B.1.351, and P.1 also suggested that some positions may be critical for modulation of S protein responses and inducing immunity escape from vaccines and different classes of monoclonal antibodies [76–79]. Together, these studies unveiled a complex balance among mutational variants in which individual modifications may act cooperatively and synergistically to enhance stability, modulate binding to the ACE2, and confer immunity resistance to neutralizing antibodies [80].

Computer simulations and protein modeling played an important role in shaping up our understanding of the dynamics and function of SARS-CoV-2 glycoproteins [81–91]. All-atom MD simulations of the full-length SARS-CoV-2 S glycoprotein embedded in the viral membrane, with a complete glycosylation profile were reported in an illuminating and insightful study by Amaro and coworkers providing the unprecedented level of details about the conformational landscapes and dynamics of S proteins in physiological environment [84]. Coarse-grained modeling of the SARS-CoV-2 virion combined results from cryo-EM and x-ray crystallography experiments together with computational predictions to build robust structural models of the SARS-CoV-2 proteins assemble a complete virion model [85]. More recent extensive MD simulations and free energy landscape mapping studies of the SARS-CoV-2 S proteins and mutants detailed conformational changes and diversity of ensembles, further supporting the notion of enhanced functional and structural plasticity of S proteins [92–97]. Our recent studies combined coarse-grained and atomistic MD simulations with coevolutionary analysis and network modeling to present evidence that the SARS-CoV-2 spike protein function as allosterically regulated machine that exploits plasticity of allosteric hotspots to fine-tune response to antibody binding [98–103]. These studies showed that examining allosteric behavior of the SARS-CoV-2 pike proteins may be useful to uncover functional mechanisms and rationalize the growing body of diverse experimental data. In particular, using atomistic-based view of allosteric communications in the SARS-CoV-2 S proteins we determined that the D614G mutation can exert its effect through allosterically induced changes on stability and communications in the residue interaction networks [104,105]. A review of computational simulation studies of the SARS-CoV-2 S proteins highlighted the synergies between experiments and simulations, outlining directions for computational biology research in understanding mechanisms of COVID-19 protein targets [106].

In this study, we employed coarse-grained (CG) simulations of closed and open states of the S-G614, S-B.1.1.7 and S-B.1.351 trimers to characterize dynamic differences in the conformational landscapes. By combining atomistic dynamic analysis with the ensemble-based local frustration analysis of the conformational states, we characterize conformational plasticity of the S variants. Through mutational scanning of protein stability changes in the conformational states, we quantify mutational adaptability of the S-G614, S-B.1.1.7 and S-B.1.351 states. The ensemble-based dynamics and energetic analysis suggests a significant conformational and mutational plasticity of the open S states for all variants. This study suggests that modulation of the energetic frustration at the inter-protomer interfaces by mutations D614G, A570D, S982A can serve as a mechanism for allosteric regulation in which the dynamic couplings between the site of mutation and the inter-protomer hinge of functional motions would modulate the inter-domain interactions, global changes in mobility and the increased stability of the open form.

## 2. Results and Discussion

### 2.1. Atomistic Modeling and Simulations Reveal Distinct Conformational Flexibility Patterns of the SARS-CoV-2 S Mutant Variants

We employed multiple CG simulations to provide a comparative analysis of the dynamic landscapes characteristic of the major functional states of the SARS-CoV-2 S trimer. While all-atom MD simulations with the explicit inclusion of the glycosylation shield could provide a rigorous assessment of conformational landscape of the SARS-CoV-2 S proteins, such direct simulations remain to be technically challenging due to the size of a complete SARS-CoV-2 S system embedded onto the membrane. We combined CG simulations with atomistic reconstruction and additional optimization by adding the glycosylated microenvironment. These cryo-EM structures of the S-G614 structures revealed structurally ordered 630 loop (residues 617 to 644) and FPPR region (residues 823 to 862) in the closed form [51] (Figure 1A,B). For the S-B.1.1.7 structures, the 630 loop is disordered in the up protomer but ordered for the down-protomer on the opposite side [61] (Figure 1C,D). The closed and open forms of the S-B.1.351 trimer are very similar to the S-G614 trimer structures [61]. Common to these structures, the all 630 loop and FPPR segment are ordered in the RBD-down form, while only FPPR and 630-loop are disordered in the 1-up protomer of the open state (Figure 1E,F). In the closed states, the NTD regions, RBD and CTD1 (residues 528-591) residues linking S1 and S2 subunits showed the increased mobility level. A comparative analysis of the dynamics profiles showed important differences in conformational flexibility of the S-B.1.1.7 and S-B.1.351 variant trimers in the closed states despite seemingly very similar structural organization and topology (Figure 2A). Notably, both S-B.1.1.7 and S-B.1.351 variants share D614G substation. The dynamics profiles suggested that other mutations in S-B.1.1.7 and S-B.1.351 may further amplify the destabilizing effect of the D614G on the stability of the closed state (Figure 2A).

**Figure 1.**
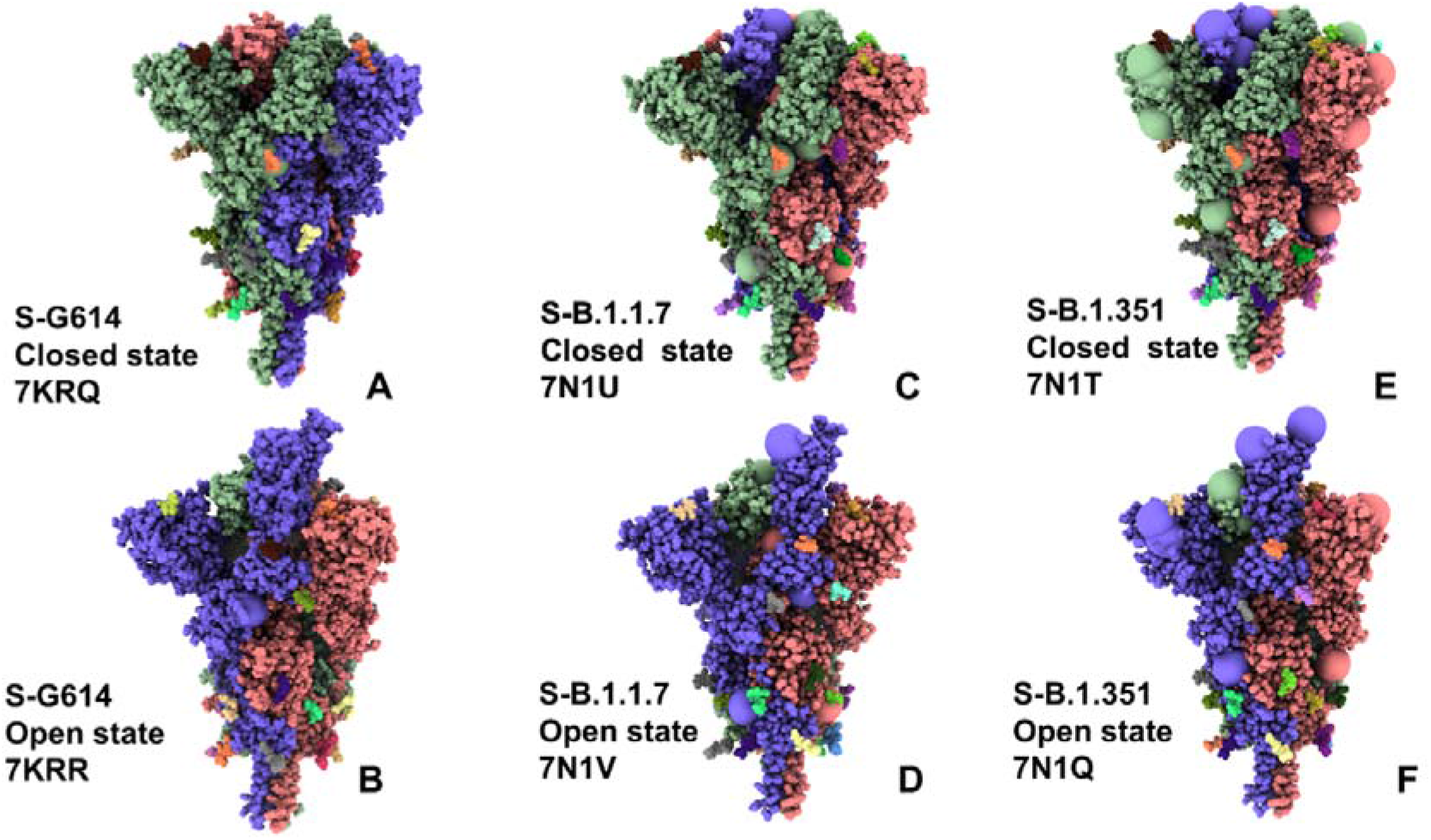
Cryo-EM structures of the SARS-CoV-2 S trimer structures used in this study. The S-G614 closed state (pdb id 7KRQ) (A) and 1 RBD-up open form (pdb id 7KRR) (B). The SARS-CoV-2 S-B.1.1.7 closed form (pdb id 7N1U) (C) and 1 RBD-up open state (pdb id 7N1V) (D). The SARS-CoV-2 S-B.1.351 closed form (pdb id 7N1T) (E) and 1 RBD-up open state (pdb id 7N1Q) (F). The structures are shown in full spheres and colored with protomers A,B,C are colored in green, red and blue. The rendering of SARS-CoV-2 S structures was done using the interactive visualization program UCSF ChimeraX [107].

**Figure 2.**
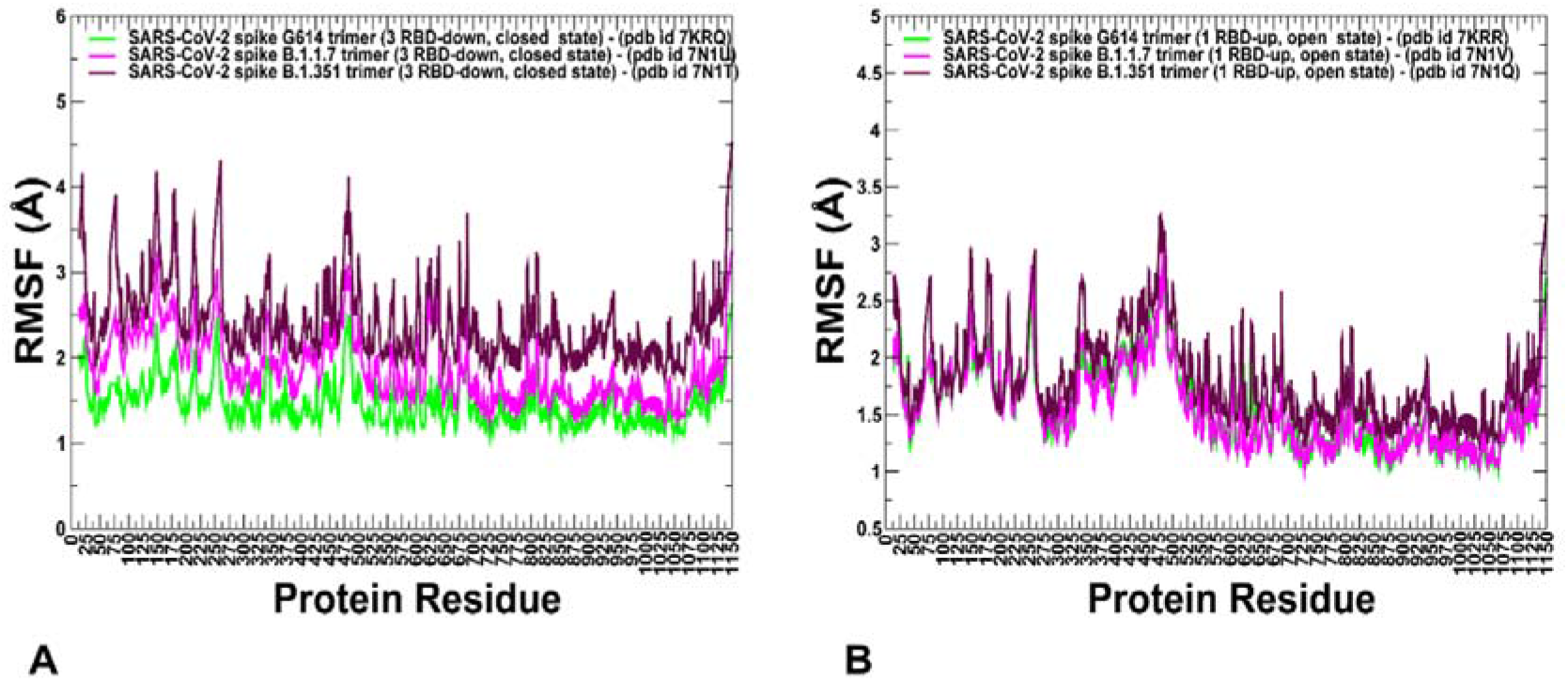
CG conformational dynamics profiles of the SARS-CoV-2 S protein variants. (A) The root mean square fluctuations (RMSF) profiles obtained from simulations of the cryo-EM structures of SARS-CoV-2 S-G614 in the closed state, pdb id 7KRQ (green lines), S-B.1.1.7 closed form, pdb id 7N1U (blue lines), and S-B.1.351 closed state, pdb id 7N1T (maroon lines). (B) The root mean square fluctuations (RMSF) profiles obtained from simulations of the cryo-EM structures of SARS-CoV-2 S-G614 in the open 1 RBD-up state, pdb id 7KRR (green lines), S-B.1.1.7 open 1 RBD-up form, pdb id 7N1V (blue lines), and S-B.1.351 open 1 RBD-up state, pdb id 7N1Q (maroon lines). The S1 regions include NTD (residues 14-306), RBD (residues 331-528), CTD1 (residues 528-591), CTD2 (residues 592-686), UH (residues 736-781), HR1 (residues 910-985), CH (residues 986-1035), CD (residues 1035-1068), HR2 (residues 1069-1163).

Interestingly, the conformational fluctuations particularly increased in the S-B.1.351 closed structure, suggesting that mutations of this variant could induce further plasticity of the closed form and facilitate transitions to the partially open form (Figure 2A). The RMSF profiles for the 1 RBD-up open states were similar for S-G614 and S-B.1.1.7 conformations, while larger fluctuations were observed in the S-B.1.351 structure (Figure 2B). Together, these findings suggested the greater variability in the functional states of the S-B. 1.351 trimer.

Structural maps of the conformational dynamics profiles illustrated the increased mobility of the closed states for the S-G614, S-B.1.1.7 and S-B.1.351 variant trimers while showing moderate flexibility of the open states (Figure 3). Of particular notice is a significant softening of the closed S-G614 and S-B.1.351 conformations. In these closed states, we detected the increased fluctuations in both S1 and S2 subunits (Figure 3 A,E). For the S-B.1.1.7 closed trimer, the fluctuations of the S2 regions were smaller and more significant flexibility was observed for the NTD regions (Figure 3C). These results suggested that the increased preferences of the S protein variants towards 1 RBD-up open conformation may be determined by the increased mobility the RBD-down closed forms. One of the key observations of the conformational dynamics analysis is the stabilization pattern in the open forms of the S protein variants (Figure 3B,D,F). In particular, we found that S-G614 and S-B.1.1.7 conformations displayed a broad stabilization in the S1 and S2 subunits but pointed to plasticity at the inter-protomer interfaces particularly near D614G site. The conformational variability in the NTD/RBD regions progressively increased in the open states from S-G614 (Figure 3B) to S-B.1.1.7 (Figure 3D) and S-B.1.351 variants (Figure 3F). Structural mapping of the conformational dynamics profiles also highlighted the increased flexibility at the S1/S2 interfaces and near inter-protomer hinges which may promote RBD movements in the up conformations for the S-B.1.1.7 and S-B.1.351 variants. These dynamic signatures of the S protein variants may imply the greater diversity of the RBD-up conformations. Interestingly, the observed plasticity of the open conformations in the S-B.1.351 variants may reflect structural heterogeneity of the RBD-up conformations revealed in cryo-EM studies [72] where the fully open state coexist with multiple substates of 1 RBD-up and 2 RBD-up conformations. Hence, our findings are consistent with the experimental studies indicating that the greater variability of the open states can allow S-B.1.351 variant to adapt RBD-up conformational responses and provide mechanisms for escape immunity from monoclonal antibodies

**Figure 3.**
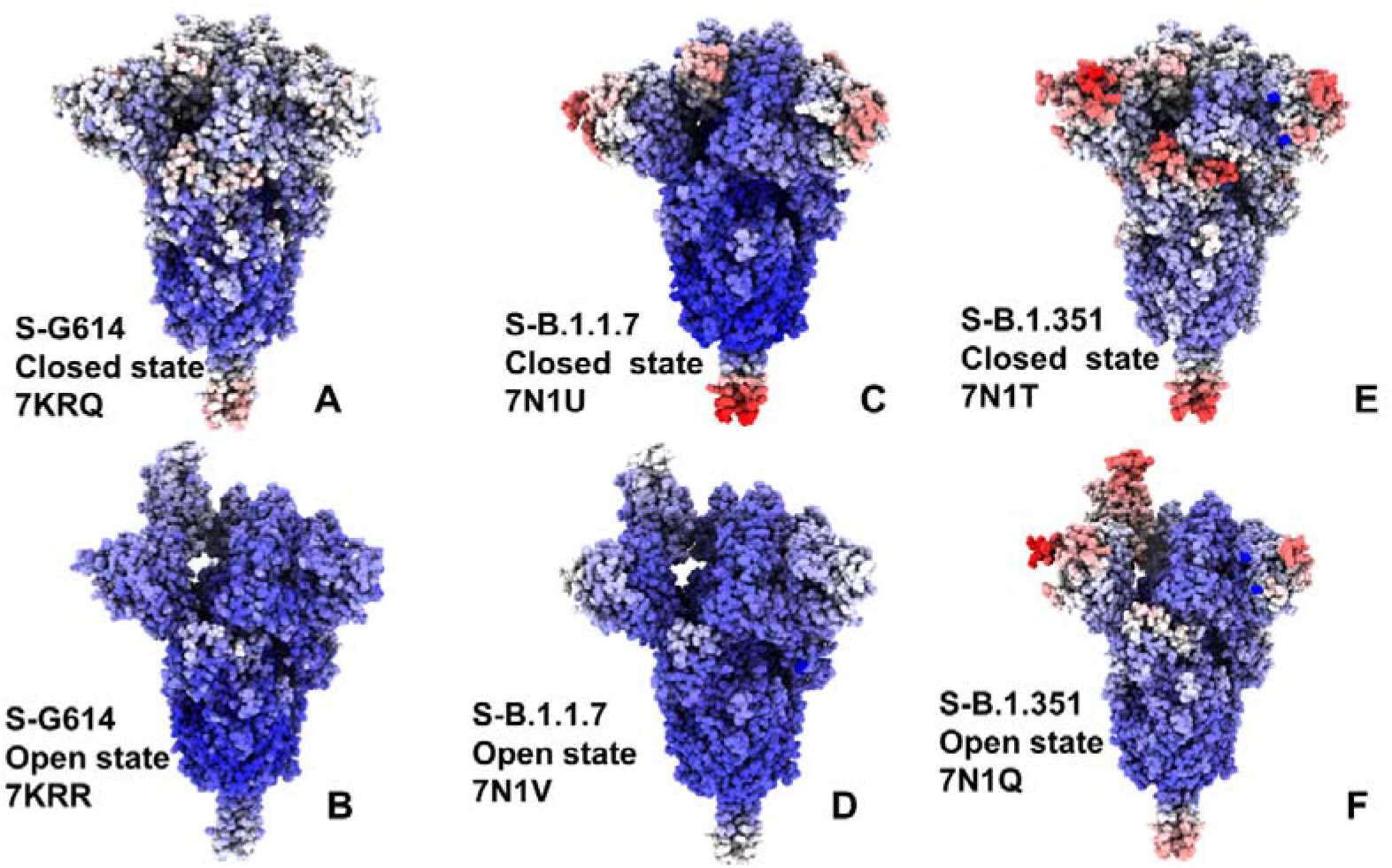
Structural maps of the conformational mobility profiles for the SARS-CoV-2 S protein variants. The dynamics map for S-G614 closed state (pdb id 7KRQ) (A) and 1 RBD-up open form (pdb id 7KRR) (B). Structural maps of the conformational mobility profiles for SARS-CoV-2 S-B.1.1.7 closed form (pdb id 7N1U) (C) and 1 RBD-up open state (pdb id 7N1V) (D). Structural maps of the conformational mobility profiles for SARS-CoV-2 S-B.1.351 closed form (pdb id 7N1T) (E) and 1 RBD-up open state (pdb id 7N1Q) (F). The structures are in sphere-based representation rendered using UCSF ChimeraX [107] with the rigidity-to-flexibility sliding scale colored from blue to red.

We also monitored relative solvent accessibility of S protein residues that was averaged over simulation trajectories (Figure 4). For the closed S-G614 ensemble, only E484 residue is solvent-exposed, while K417 and N501 are partially buried (Figure 4A). The important functional position A570 is completely buried but G614 remains only partly insulated, reflecting some plasticity at the inter-protomer interfaces near mutational site (Figure 4A). In the open S-G614 ensemble, functional residues K417, E484, N501 and G614 become moderately accessible to solvent, and only A570 position remains buried (Figure 4B). These observations reflect conformational flexibility of the RBD-up protomers and widening of the inter-protomer interfaces near G614.

**Figure 4.**
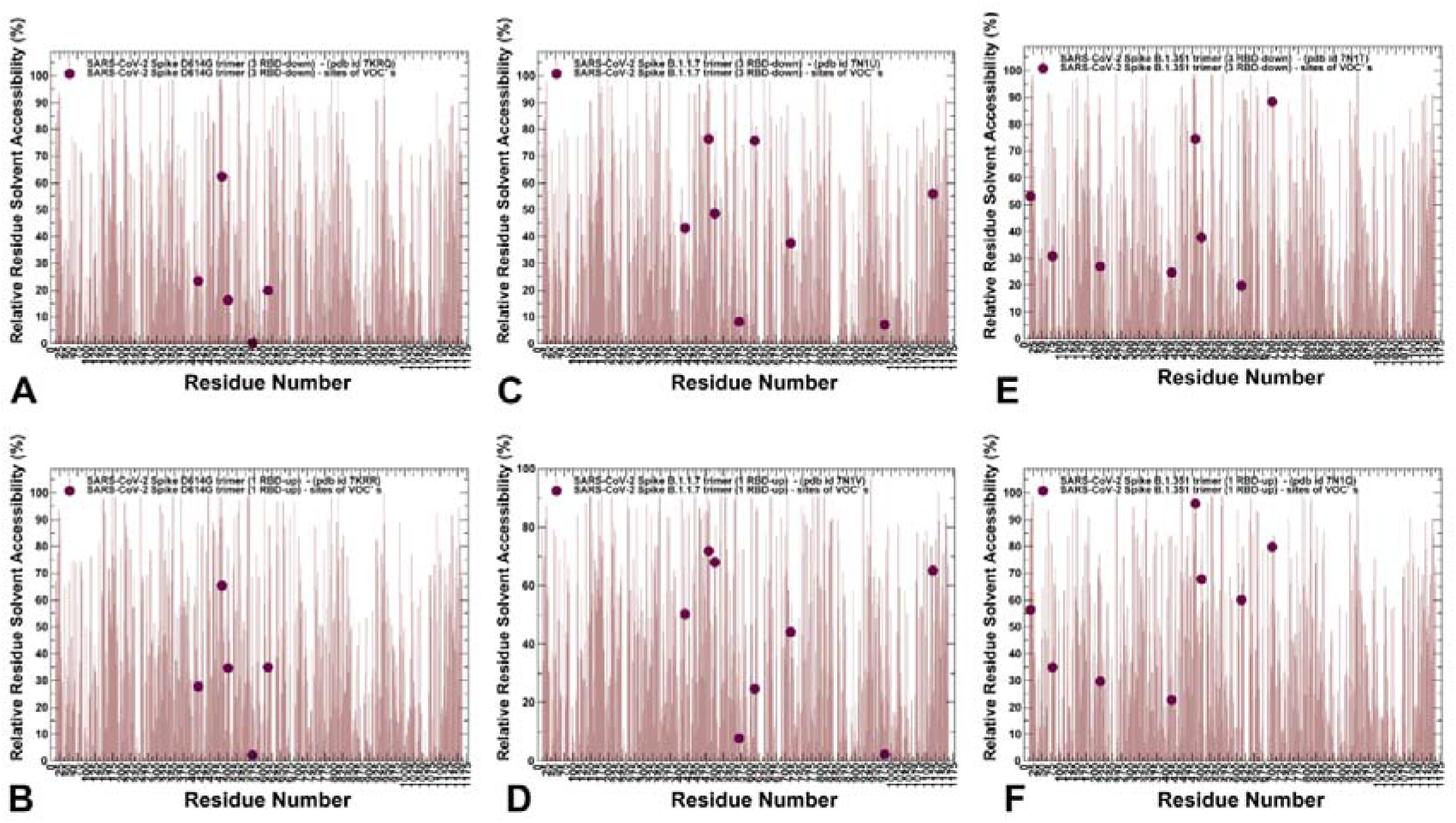
The relative residue-based solvent accessibility profiles for the SARS-CoV-2 S protein variants averaged over simulation trajectories. The average solvent accessibility profiles for the S-G614 closed state (pdb id 7KRQ) (A) and 1 RBD-up open form (pdb id 7KRR) (B). The positions of K417, E484, N501, A570, and D614G are shown in maroon-colored spheres. The average solvent accessibility profiles for SARS-CoV-2 S-B.1.1.7 closed form (pdb id 7N1U) (C) and 1 RBD-up open state (pdb id 7N1V) (D). The positions of K417, E484, N501Y, A570D, D614G, T716I, S982A, and D1118H are shown in maroon-colored spheres. The average solvent accessibility profiles forSARS-CoV-2 S-B.1.351 closed form (pdb id 7N1T) (E) and 1 RBD-up open state (pdb id 7N1Q) (F). The positions of L18F, D80A, D215G, K417N, E484K, N501Y, D614G, and A701V are shown in maroon-colored spheres.

In the S-B.1.1.7 states only A570D and S982A positions remain completely buried, while RBD sites (K417, E484, N501Y) become largely exposed and D614G position is partly exposed in both closed and open states (Figure 4C,D). The degree of solvent accessibility increased in the S-B.1.351 states (Figure 4E,F). The NTD mutational positions and RBD sites K417N, E484K, N501Y displayed an appreciable level of exposure, while D614G considerably increases its solvent exposure in the open state. These observations are consistent with the overall increase in conformational flexibility in the S-B.1.1.7 and especially S-B.1.351 variants as both closed and open forms of these variants displayed a considerable plasticity in the functional RBD regions. Interestingly, common to all S protein variants, D614G site was characterized by a moderate level of solvent accessibility, reflecting the adaptability of the S protein near the mutational site where minor structural changes allow for rearrangements in the hinge regions.

### 2.2. Functional Slow Modes of the SARS-CoV-2 S Protein Variants Reveal Role of A570D and D614G in the Hinge Regions Controlling Transitions between Open and Closed States

To identify hinge sites and characterize collective motions in the SARS-CoV-2 S-D614 and SARS-CoV-2 S-G614 structures, we performed PCA of atomistic reconstructed trajectories derived from CG simulations. Overall, the key functional signature of collective dynamics in these states are the preferences for NTD and RBD motions, suggesting that the conformational dynamics of these states can enable functional movements of RBDs to the erected, receptor-accessible conformation (Figures 5,6). The conserved hinge regions in the closed forms of the S protein variants can regulate the inter-domain movements between RBD and NTD as well as the relative motions of S1 and S2 regions. We first compared the slow mode profile averaged over the first three lowest mode for the constrained S-G614 conformations. The major hinge positions are located at F318, F592, A570, I572, Q613, G614 and Y855 residues. These sites emerged as immobilized islands in both the closed and open states. Notably, A570 and F592 hinge residues are situated near the inter-domain SD1-S2 interfaces and could act collectively as regulatory switch centers governing the population shifts between closed and open forms. For the S-B.1.1.7 and S-B.1.351 closed conformations, mutational sites A570, S982, A701, T716 are aligned with immobilized positions, while K417, E484 and N501 sites from the RBD regions belong to moving regions (Figure 5C,E). A more revealing picture emerged from the analysis of collective dynamics for the 1 RBD-up open states (Figures 5,6). In the S-G614 and S-B.1.1.7 open states, not only RBD-up undergoes functional motions but significant displacements were observed for one of the RBD-down protomers. Moreover, for the S-B.1.351 open state, functional movements were observed for all RBD’s even though the RBD-up displayed the larger magnitude of changes in the slow modes. This indicated that all S variants may promote movements of the RBD regions which may increase in the S-B.1.1.7 and especially S-B.1.351 conformations. As expected, the positions of mutational changes G614, A570, T716 and S982 in the S-B.1.1.7 states (Figure 5C,D) and positions G614 and A701 in the S-B.1.351 conformations (Figure 5E,F) corresponded to rigid sites near hinge regions, while the RBD sites K417, E484 and N501 may experience functional changes due to movements of the RBDs.

**Figure 5.**
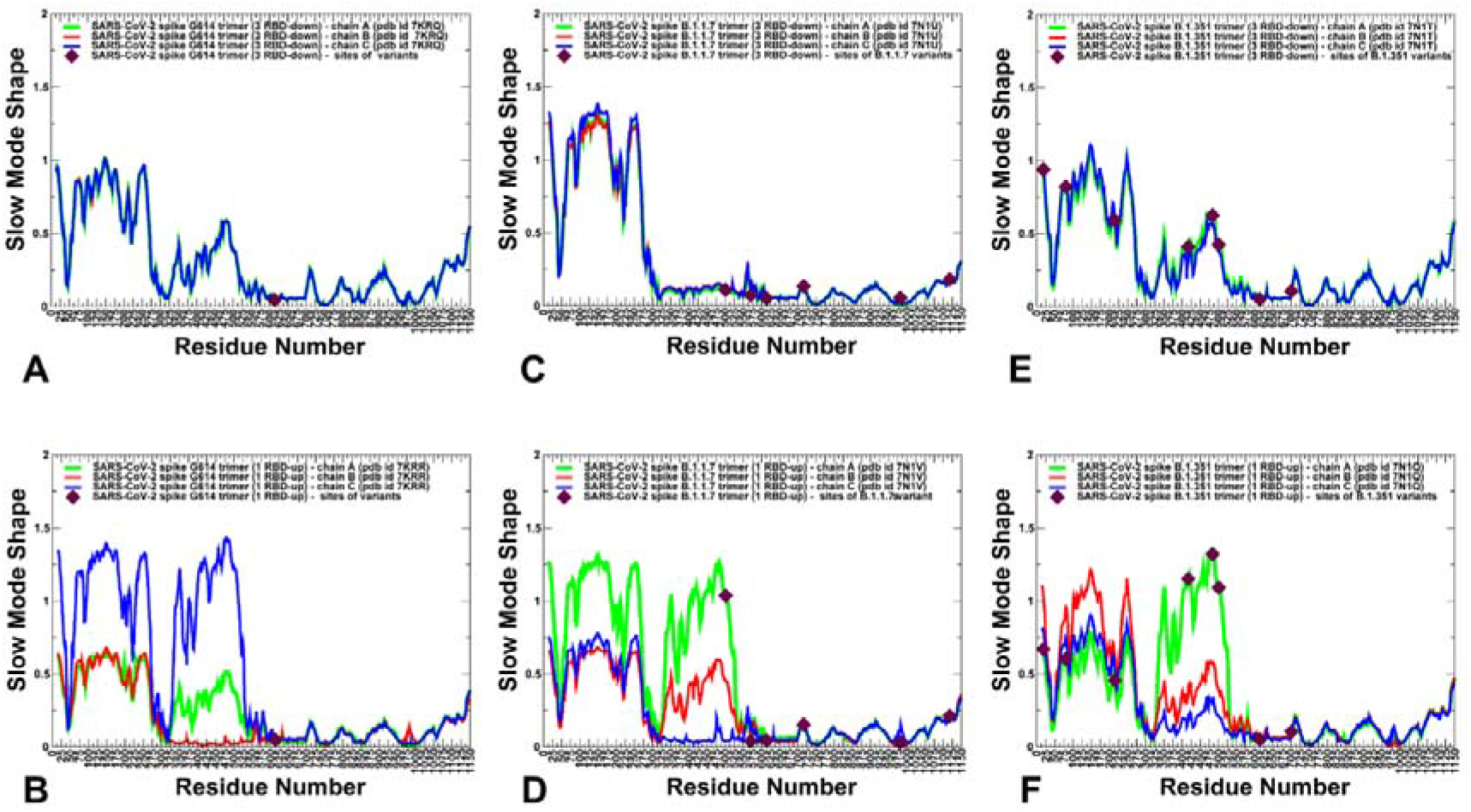
Functional dynamics of the SARS-CoV-2 S trimer structures. The essential mobility profiles are averaged over the first three major low frequency modes. The essential mobility profiles for the cryo-EM structures of SARS-CoV-2 S-G614 in the closed state, pdb id 7KRQ; (A), the open 1 RBD-up state, pdb id 7KRR (B), S-B.1.1.7 closed form, pdb id 7N1U (C), S-B.1.1.7 open 1 RBD-up form, pdb id 7N1V (D), S-B.1.351 closed state, pdb id 7N1T (E). and S-B.1.351 open 1 RBD-up state, pdb id 7N1Q (F). The profiles for protomer chains A, B and C are shown in green, red and blue lines respectively. The position of D614G is shown in maroon-colored spheres for S-G614 on panels (A,B). The positions of K417, E484, N501Y, A570D, D614G, T716I, S982A, and D1118H are shown in maroon-colored spheres for S-B.1.1.7 on panels (C,D). The positions of L18F, D80A, D215G, K417N, E484K, N501Y, D614G, and A701V are shown in maroon-colored spheres for S-B.1.351 on panels (E,F).

**Figure 6.**
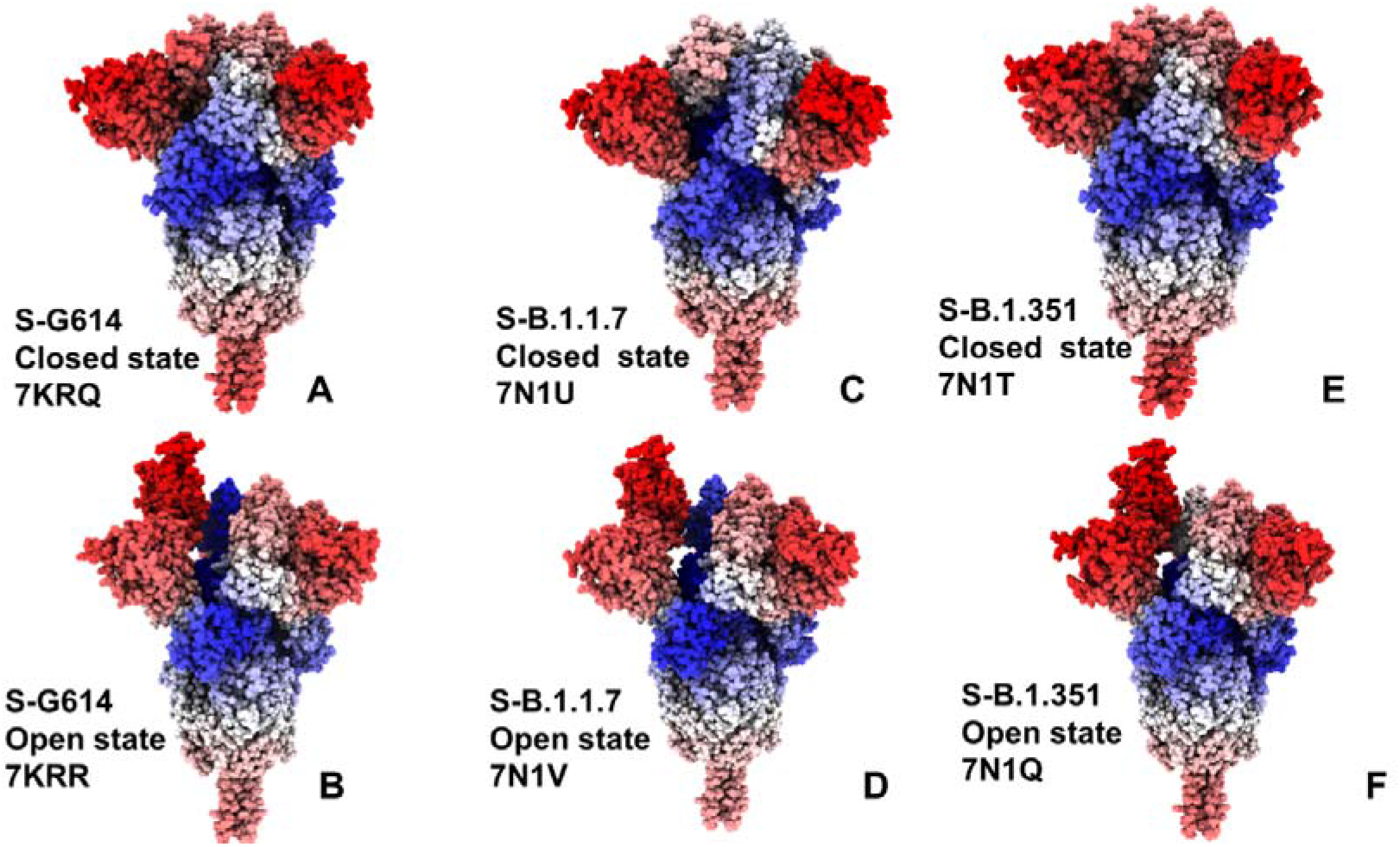
Structural maps of the essential mobility profiles for the SARS-CoV-2 S protein variants. The dynamics map for S-G614 closed state (pdb id 7KRQ) (A) and 1 RBD-up open form (pdb id 7KRR) (B). Structural maps of the conformational mobility profiles for SARS-CoV-2 S-B.1.1.7 closed form (pdb id 7N1U) (C) and 1 RBD-up open state (pdb id 7N1V) (D). Structural maps of the conformational mobility profiles for SARS-CoV-2 S-B.1.351 closed form (pdb id 7N1T) (E) and 1 RBD-up open state (pdb id 7N1Q) (F). The structures are in sphere-based representation rendered using UCSF ChimeraX [107] with the rigidity-to-flexibility sliding scale colored from blue to red.

### 2.3. Local Frustration Analysis of the SARS-CoV-2 S Conformational States: Mutational Frustration Neutrality of Variant Sites and Differential Frustration of the Closed and Open States

We employed the conformational ensembles of the S-G614, S-B.1.1.7 and S-B.1.351 trimers to estimate the average local frustration profiles of the S residues and quantify the relationship between structural plasticity and mutational frustration. This analysis is based on scanning of the conformational ensembles by local frustratometer which computes the local frustration index based on the contribution of a residue or residue pairs to the energy in a given conformation as compared to what it would contribute in decoy conformations [108–112]. The distribution of local mutational frustration showed a low relative density of highly frustrated population for residues targeted by mutations in the S trimer mutants (Figure 7). This is particularly apparent for the S-G614 conformations where K417, E484, N501, A570 and G614 sites featured a very small density of highly frustrated positions (Figure 7A). In some contrast, sites N501 and T716 in the S-B.1.1.7 conformations may be highly frustrated, while N501 and E484 may also feature an appreciable relative density of highly frustrated positions (Figure 7D). The role of frustration in the RBD regions is particularly interesting in light of recent evidence that the disordered or highly flexible regions can be critically important for mediating allostery and binding to multiple protein partners [113–117]. Notably, the population of highly frustrated conformations is relatively minor for all S conformations (Figure 7A,D,G). Importantly, we observed that sites of mutations in S-G614, S-B.1.1.7 and S-B.1.351 variants are largely associated with neutrally frustrated positions in both closed and open forms (Figure 7 B,E,H). Of notice is somewhat larger density of neutral frustration for these sites in the open states.

**Figure 7.**
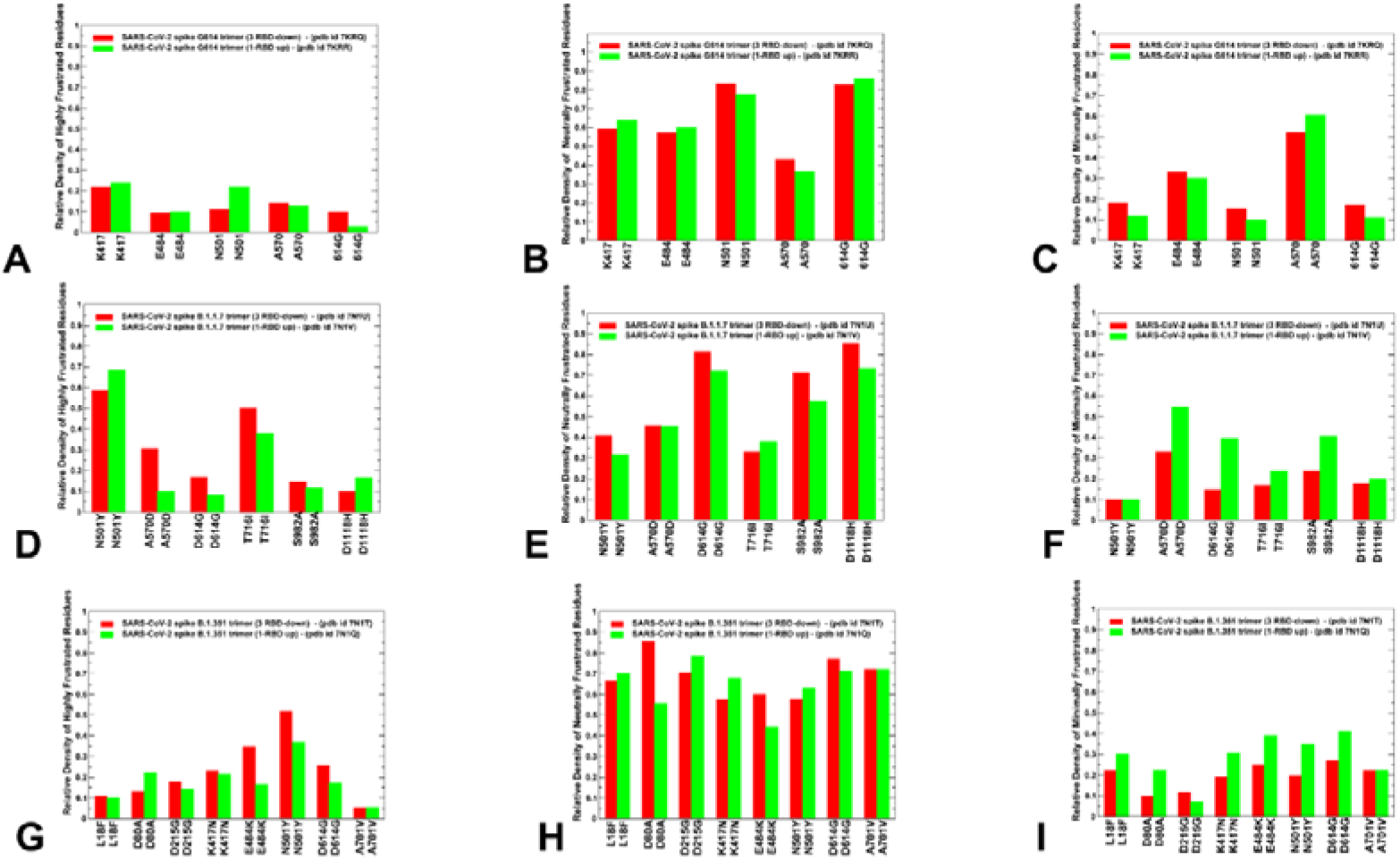
A comparison of the ensemble-averaged local mutational frustration between closed and open forms for sites of mutational variants and hinge positions of the S-G614 variant (A-C), S-B.1.1.7 variant (D-F) and S-B.1.351 variant (G-I). The relative density of highly frustrated, neutrally frustrated and minimally frustrated residues are shown. The relative density index is shown in red bars for the closed (3 RBD-down) states and in green bars for the 1 RBD-up open states.

A comparison of local mutational frustration indexes for the functional positions including sites of variants (K417, E484, N501), hinge position A570 and G614 revealed interesting shifts between the closed and open forms of S-G614 trimer (Figure 7). The high frustration density index for these positions is uniformly small and similar in both closed and open conformations. While this may be expected for hinge positions A570 and G614, it was somewhat revealing for the RBD positions, especially mobile E484 and N501 residues. The relative density of neutral frustration is pronounced for all sites, especially for the N501 and G614 positions, showing no appreciable differences between the closed and open states (Figure 7B,E). Hence, sites of circulating variants located in the flexible RBD regions showed a preponderance towards neutral mutational frustration.

There are several revealing differences in the local frustration distributions of the S-B.1.1.7 (Figure 7D,E,F) and S-B.1.351 conformations (Figure 7G,H,I). Indeed, for the S-B.1.1.7 conformations, mutational positions D614G, S982A, D1118H are mostly neutrally frustrated. At the same time, the key hinge position A570D showed a similar relative density of neutral and minimal frustration, while N501Y mutation is associated mostly with high and neutral level of frustration. This is consistent with the structural rigidity of the A570 hinge position and significant conformational plasticity of N501 site in the S-B.1.1.7 conformations. In the S-B.1.351 conformations, we observed that sites targeted by mutations in the exposed RBD and NTD regions also featured mostly neutral level of local frustration, suggesting that mutational adaptability in these positions would not perturb significantly structural stability while potentially allowing for functional plasticity during binding with the host receptor or antibodies. The relative density of minimal and high frustration for all mutational sites in the S-B.1.351 conformations is fairly low including NTD, RBD and S2 positions (Figure 7G-I). This could imply that the S-B.1.351 variant could promote moderate level of conformational variability and functional adaptability in both closed and open states.

The relationship between frustration in proteins and their function has been explored in illuminating studies by Wolynes [118]. The protein folding landscape theory established that while minimally frustrated interactions may have evolved in proteins to enable for strongly funneled landscapes, a number of functional regions and specific protein sites could have been selected to be frustrated to allow for modulation of global motions and binding adaptability with various interaction partners. In this context, our results suggested that neutral frustration patterns allow for moderately suboptimal inter-protomer and S1-S2 interactions leading to multiple states, which can be exploited to control binding with the ACE2 and to multiple partners. The results also offer an interesting rationale for the important role of A570D and D614G mutational sites in the S-B.1.1.7 and S-B.1.351 variants. These generally stable positions are involved in the inter-protomer hinge regions that enable control functional transitions between closed and open forms. Importantly, D614G is shared by all variants. We observed that in the S-G614 and S-B.1.1.7 conformations A570 an A570D positions are predominantly minimally frustrated but could display an appreciable density of neutral frustration. This minimal-to-neutral frustration level could allow for some mutational adaptability in the hinge position that retains its regulatory role in the S variants.

A slightly different distribution is characteristic of the D614G position where the predominant density of neutral frustration was observed (Figure 7). According to our analysis, there is a greater degree of mutational and conformational plasticity near the D614G position that could allow for greater variability and diversity of the open states. This may be exploited by virus to engage A570D and D614G sites in a cross-talk to modulate the thermodynamic equilibrium between the closed and open states as well as level of functional plasticity in the open substates required for allosteric function and binding. Based on our analysis, we suggest that by exploiting moderate frustration A570D and D614G substitutions in the S-B.1.1.7 and S-B.1.351 conformations may allow for the increased plasticity in the inter-protomer regions that would modulate functional motions of the RBDs without compromising stability of the S1-S2 interfaces.

The generally prevailing pattern of frustration neutrality for sites targeted by mutations across all studied VOC’s and dynamic contributions of high and neutral frustration in the RBD sites E484 and N501 are important findings of this analysis. The highly frustrated interactions may in principle conflict with the robust folding of the domain and could reflect evolutionary constraints other than folding. This may explain the specific local frustration pattern of the A570 and D614 positions that combined could provide moderate adaptability of the inter-protomer regions. On the other hand, local frustration is often an important driver of allosteric changes. Our results indicated that high-to-neutral frustration level is characteristic of the RBD positions targeted by mutations, especially N501. The relatively high frustration in these positions may allow for discrete set of configurations involving local motions of the frustrated residues. Combined with modulation of the inter-protomer hinge regions by A570D and D614G positions, the local frustration in the RBD mutational sites could drive dynamical transitions between closed and open states accompanied by local adjustments of the RBD residues.

### 2.3. Mutational Scanning of Protein Stability the SARS-CoV-2 S-614 Conformational States Reveals Energetic Effects of the D614G Mutation

We employed the equilibrium ensembles generated by simulations of the SARS-CoV-2 protein structures to perform mutational scanning of the spike protein residues and mutational sensitivity analysis of the S-B.1.1.7 (Figure 8) and S-B.1.351 mutant proteins (Figure 9). The protein stability ΔΔG changes were computed by averaging the results of computations over 1,000 samples obtained from simulation trajectories. Using mutational cartography, we first examined the pattern of free energy changes for the S-B.1.1.7 closed and open states (Figure 8). Interestingly, it appeared that mutations N501Y, A570D and D614G produced larger destabilization changes in the open state (Figure 8B). This is consistent with the dynamics analysis indicating that S-B.1.1.7 mutations can significantly destabilize the closed state and moderately stabilize the open state (Figures 2,3). Importantly, these differences were found in the key positions that collectively responsible for modulation of the inter-protomer interactions and positioning of the RBD regions. Mutations in other functional RBD positions (K417, E484) as well as in T716, S982 and D1118 positions resulted in the minor energy changes in both closed (Figure 8A) and open S-B.1.1.7 conformations (Figure 8B). S982A substitution in the S-B.1.1.7 conformations abolished hydrogen bonding between central helices of the S2 domain and G545 in the CTD1 region [70]. We found that mutations of A982 to other positions in the S-B.1.1.7 states resulted in moderate and often stabilizing changes. Hence, this position is a relatively soft residue where mutations can be tolerated without significant impact on the protein stability. The mutational cartography analysis revealed that A570D and D614G mutational positions in the S-B.1.1.7 conformations are the most sensitive to modifications which result in more significant destabilizing changes (Figure 8). Importantly, the effect of mutations in these positions on the protein stability is greater in the open state (Figure 8B). The S-B.1.1.7 open state featured a salt bridge involving interactions of A570D with K854 of the other protomer [70]. In the closed S-B.1.1.7 states A570D can form the inter-protomer interactions with K964 and N856 that together comprise an important hinge cluster. Functional dynamics analysis of slow modes confirmed the role of A570D as a potential regulatory switch that controls RBD movements. Although modifications of A570D are generally destabilizing, the range of free energy changes associated with this position suggested a moderate level of residual energetic frustration and suboptimal interactions (Figure 8). These findings are consistent with a series of structural studies showing a moderate degree of conformational heterogeneity in the interprotomer interactions formed by A570D in different substates of the S-B.1.1.7 protein [70,71]. Interestingly, substitutions in the D614G position are more destabilizing in both closed and open S-B.1.1.7 states (Figure 8), pointing to dynamic rearrangements near D614G position.

**Figure 8.**
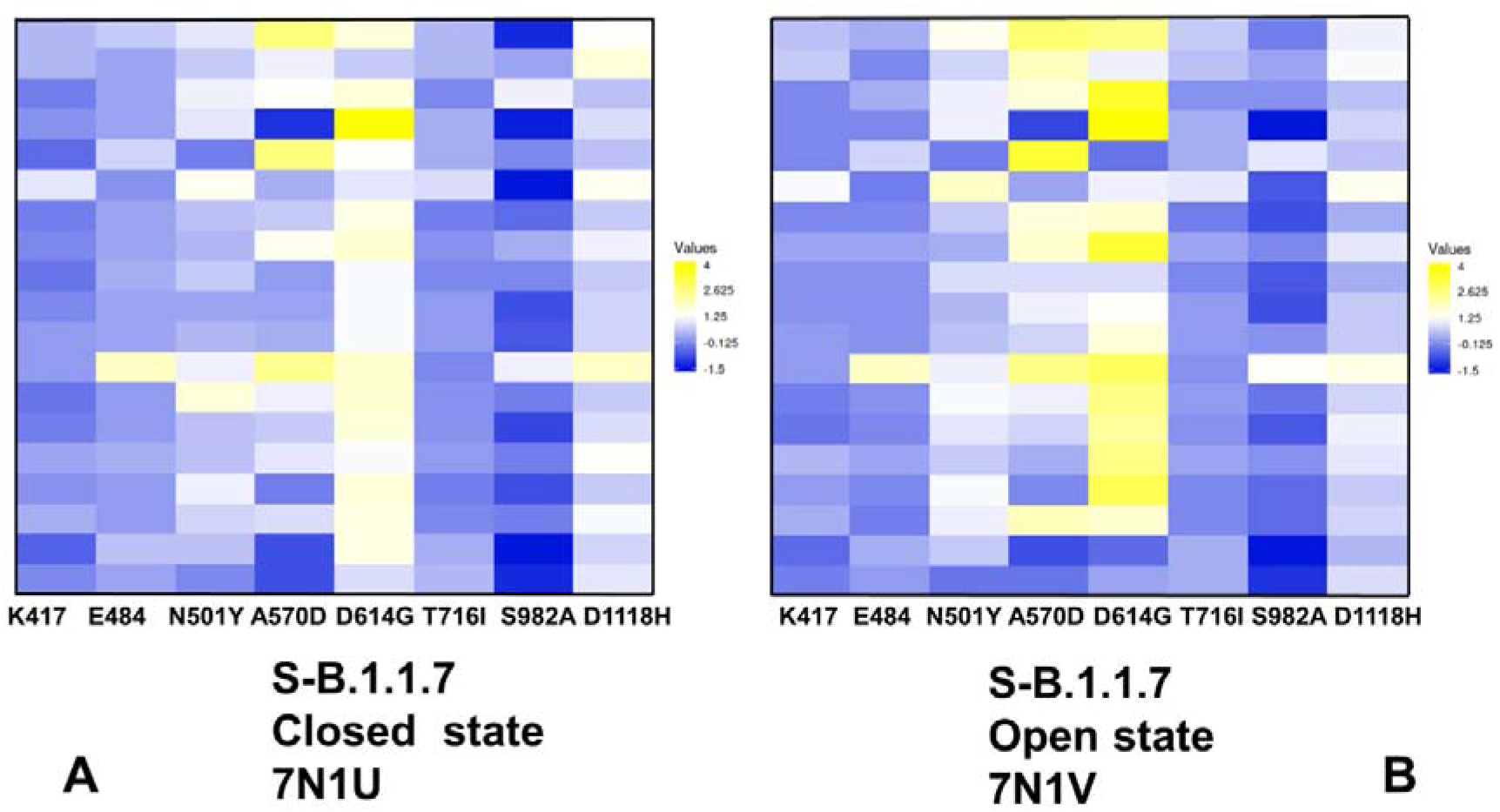
Ensemble-based mutational profiling of the SARS-CoV-2 S-B.1.1.7 protein stability. The mutational scanning heatmaps are shown for the closed state (A) and open state (B). The heatmaps show the computed binding free energy changes for 19 single mutations on the sites of variants. The squares on the heatmap are colored using a 3-colored scale – from blue to white and to yellow, with yellow indicating the largest unfavorable effect on stability. The standard errors of the mean for binding free energy changes were based on 5 independent CG trajectories for each of the S-B.1.1.7 states and different number of selected samples from a given trajectory (500, 1,000 and 2,000 samples) are ~ 0.11-0.24 kcal/mol using averages over different trajectories and ≤ 0.15 kcal/mol from computations based on different number of samples from a single trajectory.

**Figure 9.**
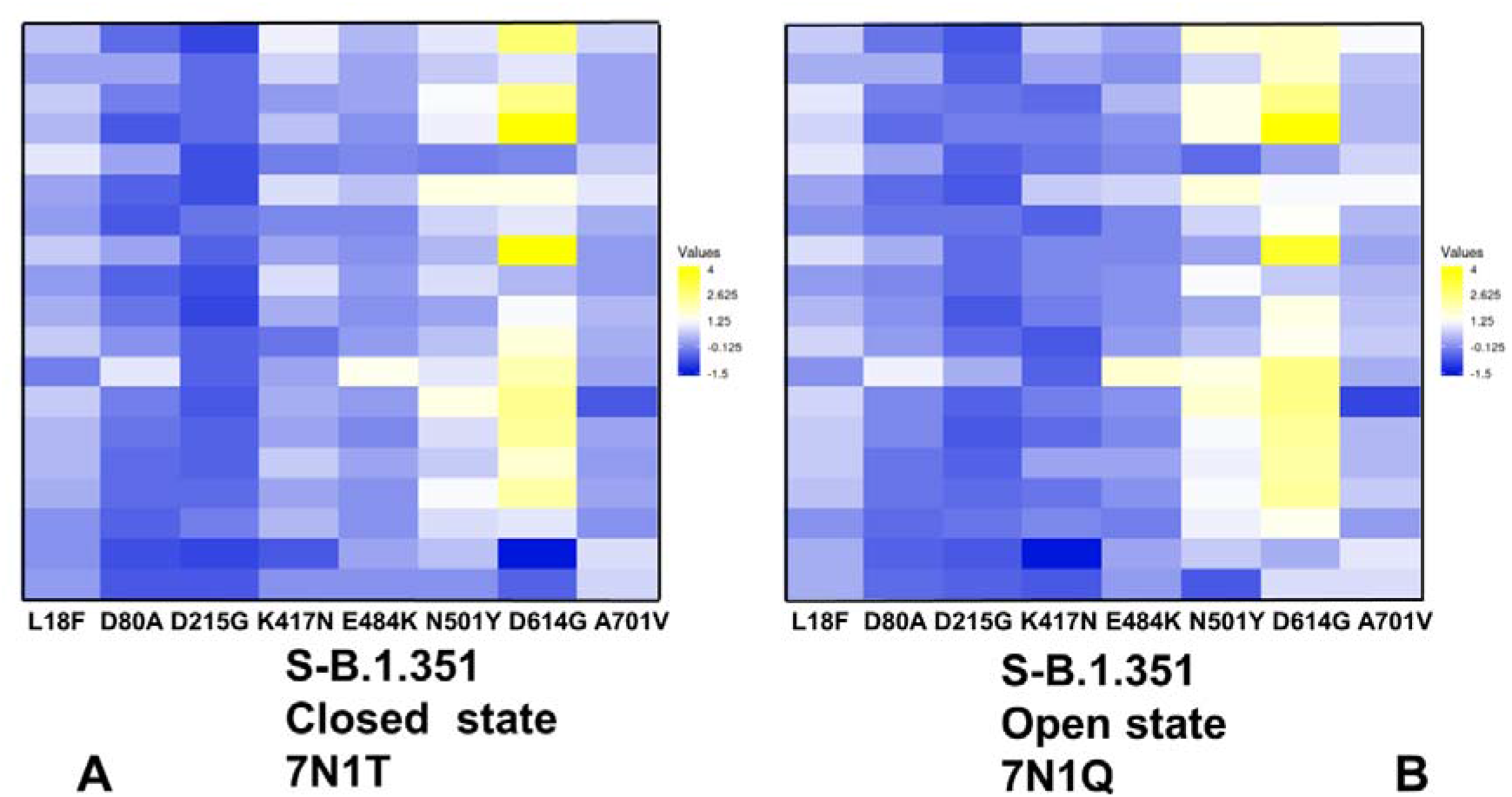
Ensemble-based mutational profiling of the SARS-CoV-2 S-B.1.351 protein stability. The mutational scanning heatmaps are shown for the closed state (A) and open state (B). The heatmaps show the computed binding free energy changes for 19 single mutations on the sites of variants. The squares on the heatmap are colored using a 3-colored scale – from blue to white and to yellow, with yellow indicating the largest unfavorable effect on stability. The standard errors of the mean for binding free energy changes were based on 5 independent CG trajectories for each of the S-B.1.351 states and different number of selected samples from a given trajectory (500, 1,000 and 2,000 samples) are ~ 0.18-0.27 kcal/mol using averages over different trajectories and ≤ 0.17 kcal/mol from computations based on different number of samples from a single trajectory.

Together, A570D and D614G sites are involved in the inter-protomer interactions in the S-B.1.1.7 states and contribute to the hinge clusters that modulate RBD motions. The results of mutational scanning are supportive of the local frustration analysis that displayed neutral-to minimal frustration densities for A570D and D614G positions in the S-B.1.1.7 state. We suggest that neutral-to-minimal level of energetic frustration and moderate conformational plasticity in these regions and near the inter-domain interfaces could allow for emergence of mutational variants in sites responsible for allosteric modulation of conformational transitions between closed and open S states.

Mutational scanning maps for variant positions in the S-B.1.351 conformations showed similar and moderate stabilization changes for the NTD variant L18F, D80A, and D215G. It is evident that many substitutions in G215 position can be in fact energetically favorable for the protein stability (Figure 9). This pattern for the NTD mutational sites is shared in the closed and open states, suggesting that these variants could promote destabilization of the S-B.1.351 conformations and increase mobility and conformational variability of the NTDs. Moderate free energy changes are also seen for K417N and E484K sites in the closed and open states. Interestingly, there is a clear difference in the mutational map for the N501Y position (Figure 9). While variations in this position are only moderately destabilizing in the closed state, the destabilization is increased in the open S-B.1.351 conformation. Hence, N501Y may become less frustrated in the open state and provide for more optimal interactions of the RBDs. In both states of S-B.1.351 variant, we observed appreciable destabilization changes induced by modifications in the D614G position (Figure 9). This is consistent with the increased density of minimal frustration for this position in the open S-B1.351 state.

Importantly, the results showed a significant difference between S-B.1.1.7 and S-B.1.351 structures in the mutational profiles for K417/E484/N501 positions (Figures 8,9). We found that these K417N, E484K and N501Y sites can afford greater mutational tolerance and conformational plasticity in the S-B.1.351 conformations. This suggested that the level of mutational and conformational plasticity can progressively increase from S-G614 to S-B.1.1.7 and S-B.1.351 variants. The results also suggested an interesting interplay between mutation-induced protein stability and local frustration patterns. It is interesting to examine these relationships in the context of allosteric regulation models centered on the energetic frustration that could emerge at the inter-domain interfaces [113–117]. In these models, activation or repression functions may be encoded in the conformational ensemble and reveal through frustration-based allosteric regulation of the inter-domain interactions. The energetic scanning and local frustration analyses revealed the inherent mutational plasticity of the sites targeted by circulating variants. Mutational positions that are involved in hinge clusters are characterized by some degree of energetic frustration, thereby allowing for allosteric couplings and modulation of the RBD motions leading to the greater diversity of RBD-exposed conformations. These results suggested that S protein variants may uniquely exploit the intrinsic conformational and mutational plasticity of the S proteins that is broadly distributed and is characteristic of not only RBD regions but also present in the inter-protomer and inter-domain regions. We argue that S-B.1.1.7 and S-B.1.351 variants may leverage this plasticity to adopt a mechanism of frustration-based allosteric modulation at the inter-protomer interfaces to differentially control binding with the host cell receptor ACE2 and interacting proteins.

## 3. Materials and Methods

### 3.1 Structure Preparation and Analysis

All structures were obtained from the Protein Data Bank [119,120]. During structure preparation stage, protein residues in the crystal structures were inspected for missing residues and protons. Hydrogen atoms and missing residues were initially added and assigned according to the WHATIF program web interface [121,122]. The structures were further pre-processed through the Protein Preparation Wizard (Schrödinger, LLC, New York, NY) and included the check of bond order, assignment and adjustment of ionization states, formation of disulphide bonds, removal of crystallographic water molecules and co-factors, capping of the termini, assignment of partial charges, and addition of possible missing atoms and side chains that were not assigned in the initial processing with the WHATIF program. The missing loops in the studied cryo-EM structures of the SARS-CoV-2 S protein were reconstructed and optimized using template-based loop prediction approaches ModLoop [123], ArchPRED server [124] and further confirmed by FALC (Fragment Assembly and Loop Closure) program [125]. The side chain rotamers were refined and optimized by SCWRL4 tool [126]. The conformational ensembles were also subjected to all-atom reconstruction using PULCHRA method [127] and CG2AA tool [128] to produce atomistic models of simulation trajectories. The protein structures were then optimized using atomic-level energy minimization with a composite physics and knowledge-based force fields as implemented in the 3Drefine method [129]. The atomistic structures from simulation trajectories were further elaborated by adding N-acetyl glycosamine (NAG) glycan residues and optimized. The glycosylated microenvironment for atomistic models of the simulation trajectories was mimicked by using the structurally resolved glycan conformations for 22 most occupied N-glycans [130,131] in each as determined in the cryo-EM structures of the SARS-CoV-2 spike S trimer in the closed state (K986P/V987P,) (pdb id 6VXX) and open state (pdb id 6VYB), and the cryo-EM structure SARS-CoV-2 spike trimer (K986P/V987P) in the open state (pdb id 6VSB).

### 3.2 Coarse-Grained Simulations

Coarse-grained (CG) models are computationally effective approaches for simulations of large systems over long timescales. We employed CABS-flex approach that efficiently combines a high-resolution coarse-grained model and efficient search protocol capable of accurately reproducing all-atom MD simulation trajectories and dynamic profiles of large biomolecules on a long time scale [132–138]. In this high-resolution model, the amino acid residues are represented by Cα, Cβ, the center of mass of side chains and another pseudoatom placed in the center of the Cα-Cα pseudo-bond. In this model, the amino acid residues are represented by Cα, Cβ, the center of mass of side chains and the center of the Cα-Cα pseudo-bond. The CABS-flex approach implemented as a Python 2.7 object-oriented standalone package was used in this study to allow for robust conformational sampling proven to accurately recapitulate all-atom MD simulation trajectories of proteins on a long time scale [136–138]. Conformational sampling in the CABS-flex approach is conducted with the aid of Monte Carlo replica-exchange dynamics and involves local moves of individual amino acids in the protein structure and global moves of small fragments [136]. The default settings were used in which soft native-like restraints are imposed only on pairs of residues fulfilling the following conditions : the distance between their C^α^ atoms was smaller than 8 Å, and both residues belong to the same secondary structure elements. A total of 1,000 independent CG-CABS simulations were performed for each of the studied systems. In each simulation, the total number of cycles was set to 10,000 and the number of cycles between trajectory frames was 100. MODELLER-based reconstruction of simulation trajectories to all-atom representation provided by the CABS-flex package was employed to produce atomistic models of the equilibrium ensembles for studied systems [136]. We also performed principal component analysis (PCA) of reconstructed trajectories derived from CABS-CG simulations using the CARMA package [139].

### 3.3 Mutational Scanning and Sensitivity Analysis

To compute protein stability changes in the SARS-CoV-2 S structures, we conducted a systematic alanine scanning of protein residues in the SARS-CoV-2 trimer mutants as well as mutational sensitivity analysis at the mutational site for both SARS-CoV-2 S-D614 and SARS-CoV-2 S-G614 structures. Two different approaches were used. Alanine scanning of protein residues was performed using FoldX approach [140–145]. and BeAtMuSiC approach [146–148]. If a free energy change between a mutant and the wild type (WT) proteins ΔΔG= ΔG (MT)-ΔG (WT) > 0, the mutation is destabilizing, while when ΔΔG <0 the respective mutation is stabilizing. BeAtMuSiC approach is based on statistical potentials describing the pairwise inter-residue distances, backbone torsion angles and solvent accessibilities, and considers the effect of the mutation on the strength of the interactions at the interface and on the overall stability of the complex.^121-123^ We computed the ensemble-averaged binding free energy changes using equilibrium samples from atomistic trajectories. The binding free energy changes were computed by averaging the results over 100 trajectory samples for each of the studied systems.

## 4. Conclusions

In this work, we combined molecular simulations and collective dynamics analysis with the ensemble-based frustration analysis to characterize conformational plasticity and functional adaptability of the closed and open states for the S-G614, S-B.1.1.7 and S-B.1.351 variant. The conformational dynamics analysis revealed progressively increased mobility of the S-B.1.1.7 and S-B.1.351 variant in the open states. Collective dynamics of the S protein variants confirmed the critical regulatory role of the A570D and D614G positions acting as components of hinge clusters controlling the transitions between closed and open forms. This analysis suggested that in the S-B.1.1.7 and S-B.1.351 variants all RBDs can experience significant functional movements including the RBD-down conformations. We found that mutations in the S variants may promote movements of the RBD regions which may increase in the S-B.1.1.7 and especially S-B.1.351 conformations. The local frustration analysis showed a prevailing pattern of frustration neutrality for sites targeted by mutations across all studied variant and dynamic contributions of high and neutral frustration in the RBD sites E484 and N501 positions. A strong preference of the mutational sites towards neutrally frustrated environment may allow for moderately suboptimal inter-protomer interactions and subtle control of S binding with the ACE2 and to multiple partners. Mutational scanning of protein stability revealed that level of mutational and energetic plasticity can progressively increase from S-G614 to the S-B.1.1.7 and S-B.1.351 variants. The results indicated that K417N, E484K and N501Y sites can afford greater mutational tolerance and conformational plasticity in the S-B.1.351 conformations. The results also suggested an interesting interplay between mutation-induced protein stability, local frustration and allosteric modulation of the S protein dynamics. We found that the A570D and D614G mutations may introduce a moderate level of energetic frustration and conformational plasticity near the inter-protomer and inter-domain interfaces. Coupled with flexibility of more frustrated RBD mutational sites, this could allow for allosteric couplings between the hinge clusters and RBD regions. Through this frustration-driven allosteric mechanism, mutational variants can impose allosteric control over functional movements and conformational diversity of the RBD regions. This landscape-based mechanism of mutation-induced energetic frustration in the S protein may result in the greater adaptability and the emergence of multiple conformational substates in the open form.

## Author Contributions

Conceptualization, G.V.; methodology, G.V.; software, G.V.; validation, G.V.; formal analysis, G.V.; investigation, G.V.; resources, G.V.; data curation, G.V.; writing—original draft preparation, G.V.; writing—review and editing, G.V.; visualization, G.V.; supervision, G.V.; project administration, G.V.; funding acquisition, G.V. All authors have read and agreed to the published version of the manuscript.

## Acknowledgments

The author thanks Schmid College of Science and Technology, Chapman University for providing computing resources at the Keck Center for Science and Engineering

